# Microhabitats shape diversity-productivity relationships in freshwater bacterial communities

**DOI:** 10.1101/231688

**Authors:** Marian L. Schmidt, Bopaiah A. Biddanda, Anthony D. Weinke, Edna Chiang, Fallon Januska, Ruben Props, Vincent J. Denef

## Abstract

Eukaryotic communities commonly display a positive relationship between biodiversity and ecosystem function (BEF) but the results have been mixed when assessed in bacterial communities. Habitat heterogeneity, a factor in eukaryotic BEFs, may explain these variable observations but it has not been thoroughly evaluated in bacterial communities. Here, we examined the impact of habitat on the relationship between diversity assessed based on richness, evenness, or phylogenetic diversity, and heterotrophic productivity. We sampled co-occurring free-living (more homogenous) and particle-associated (more heterogeneous) bacterial habitats in a freshwater, estuarine lake. Diversity measures, and not environmental variables, were the best predictors of particle-associated heterotrophic production. There was a strong, positive, linear relationship between particle-associated bacterial richness and heterotrophic productivity that strengthened with evenness. There were no observable BEF trends in free-living bacterial communities. Across both habitats, communities with more phylogenetically related taxa had higher per-capita heterotrophic production than communities of phylogenetically distantly related taxa. Our findings show that heterotrophic bacterial productivity is positively correlated with evenness and richness, negatively with phylogenetic diversity, and that BEF relationships are contingent on microhabitats. Our work adds to the understanding of the highly distinct contributions to community diversity and ecosystem functioning contributed by bacteria in free-living and particle-associated aquatic habitats.

## Introduction

Our planet is currently experiencing an extreme species extinction event (Thomas et al., 2004; Wake & Vredenburg, 2008, Ferrier et al., 2016). Concern about such declines in biodiversity has resulted in hundreds of studies evaluating the relationship between biodiversity and ecosystem function (BEF), with an even larger biodiversity effect in natural ecosystems compared to controlled experiments (Duffy et al., 2017). BEF relationships focusing on the number of species (*i.e.* species richness) as the biodiversity value are generally positive and asymptotic and thus biodiversity loss causes a small change in ecosystem function at first and then, at some tipping point, a dramatic decrease in function (Cardinale et al., 2012; Tilman et al., 2014). While the focus of local and global species loss is typically on eukaryotic organisms, bacterial species numbers have also been found to be decreasing at local scales within the human gut (Blaser, 2014) and terrestrial ecosystems (Singh et al., 2014). Of particular concern is the loss of the number of bacterial guilds responsible for key geochemical transformations, such as methane oxidation (Levine et al., 2011) that controls rates of methane emissions.

An extrapolation from eukaryotic relationships would predict there to be no richness-ecosystem function relationships for bacterial communities because they are generally composed of an order of magnitude more taxa than the communities in most eukaryotic BEF studies. Several studies have indicated no relationship between species richness with broad functional processes, such as heterotrophic respiration or biomass production, that are performed by many taxa (see figure 5 in Levine et al., 2011; Langenheder et al., 2006; Delgado-Baquerizo et al., 2016). Yet, other studies on broad processes such as denitrification (Philippot et al., 2013) and on narrow metabolic processes that are catalyzed by few bacterial taxa, such as methanotrophy (Levine et al., 2011), and the degradation of triclosan and microcystin (Delgado-Baquerizo et al., 2016) did find evidence of bacterial richness and ecosystem function relationships. In addition to the number of species, there are several other components of biodiversity including functional trait or gene richness (*i.e.* abundance-unweighted) (Reich et al., 2004; Flynn et al., 2011; Evans et al., 2017), species or functional dominance or evenness (*i.e.* abundance-weighted) (Wilsey & Potvin, 2000; Wilsey & Polley, 2004; Kirwan et al., 2007; Wittebolle et al., 2009), and phylogenetic diversity metrics (Flynn et al., 2011). Plant community evenness has been shown to impact plant productivity more than richness (Wilsey & Potvin, 2000; Wilsey & Polley, 2004; Kirwan et al., 2007) and the initial evenness of microbial microcosms has been found to be a key factor in functional stability, even under the selective stressors of temperature and salt stress (Wittebolle et al., 2009)

Phylogenetic relatedness, while having received less attention than richness and evenness, may also influence BEF relationships. Indeed, some studies show relationships across different ecosystems between phylogenetic diversity and ecosystem functions (Cadotte et al., 2008; Jiang et al., 2010). However, research with freshwater green algae (Fritschie et al., 2014; Venail et al., 2014) did not find this relationship. A more recent study found the opposite result by showing that closely related green algal species had weaker competition and more facilitation than distantly related species, thus resulting in higher productivity (Narwani et al., 2017). While relationships between phylogenetic relatedness among community members and ecosystem function have been assessed in bacterial systems (Tan et al., 2012; Galand et al., 2015; Roger et al., 2016), much work has focused on low-diversity, experimentally-assembled communities with bacteria that can be grown in culture. It is worthwhile to expand these findings to communities with richness levels typically found in natural communities.

The nature of BEF relationships and the mechanism(s) that underpins them may depend on habitat structure or heterogeneity. Increasing habitat heterogeneity has been found to enhance the strength of BEF relationships (Tylianakis et al. 2008), presumably due to a greater role for niche complementarity effects in heterogeneous environments (Cardinale 2011). While habitat heterogeneity contributes to increased diversity within bacterial populations and communities (Zhou et al., 2008; Shade et al., 2008), the influence of habitat heterogeneity on BEF relationships remains unknown for bacterial systems.

In this study, we hypothesized that bacterial diversity would be positively correlated with bacterial heterotrophic production, and that this relationship would be stronger in more heterogeneous environments. We simultaneously surveyed free-living and particle-associated surface water bacterial communities. These habitats have been extensively studied for their ability to sustain distinct bacterial communities and ecosystem processes (Crump et al., 1999; Bižić-Ionescu et al., 2014; Mohit et al., 2014; Schmidt et al., 2016). In addition, studies on model aggregates had 3-fold higher bacterial protein production and two orders of magnitude higher protease activity (Grossart et al., 2007), indicating particles can be an important place for microbial activity in aquatic ecosystems. Particulate matter comprises a variety of types and sizes of particles with some particles also harboring physicochemical gradients (Simon et al., 2002), and hence represents a more heterogeneous habitat than the surrounding water. We tested BEF relationships using a variety of diversity metrics including observed richness, evenness, and phylogenetic diversity. We focused on heterotrophic bacterial production as our measure of ecosystem function, as it is a key process affecting freshwater bacterial growth that in turn fuels the eukaryotic food web (Cotner & Biddanda, 2002).

## Methods

### Lake sampling and sample processing

Surface water samples were collected at 1 meter depth from 4 long-term sampling stations (Steinman et al., 2008) in mesotrophic Muskegon Lake (**Figure S1**), which is a freshwater estuarine lake connecting the Muskegon River and Lake Michigan. These stations included the mouth of the Muskegon River (43.250133,-86.2557), the channel to Bear Lake (43.238717,-86.299283; a hypereutrophic lake), channel to Lake Michigan (43.2333,-86.3229; oligotrophic lake), and the deepest basin of Muskegon Lake (43.223917,-86.2972; max depth = 24 m).

Samples were collected during the morning to early afternoon of 3 days in 2015 (May 12, July 21, & September 30) aboard the R/V W.G. *Jackson*. All water samples were collected with vertical Van Dorn samplers. Additionally, a vertical profile of temperature (T), pH, specific conductivity (SPC), oxidation-reduction potential (ORP), chlorophyll (Chl*a*), total dissolved solids (TDS), and dissolved oxygen (DO) was constructed at each station to characterize the water column using a calibrated YSI 6600 V2-4 multiparameter water quality sonde (Yellow Springs Instruments Inc.). Total Kjeldahl nitrogen (TKN), ammonia (NH3), total phosphorus (TP), and alkalinity (Alk) were processed from whole water while nitrate (NO3), phosphate (PO4), and chloride (Cl-) were hand filtered using a 60 mL syringe fitted with Sweeny filter holder with a 13 mm diameter 0.45 µm pore size nitrocellulose filters (Millipore) and were determined by standard wet chemistry methods in the laboratory (EPA, 1993).

### Bacterial abundance by epifluorescence microscopy

Lake surface water samples were processed within 2-6 hours of their collection for determination of heterotrophic bacterial abundance. Unfiltered lake water samples (5 mL) were preserved with 2% formalin and 1 mL subsamples were stained with acridine orange stain and filtered onto black 25 mm 0.2 μm pore size polycarbonate filters (Millipore) at a maximum pressure of 0.1 Bar or 1.5 PSI. Prepared slides were stored frozen until manual enumeration by standard epifluorescence microscopy at 1000x magnification under blue light excitation (Hobbie et al. 1977). Bacteria within the field of view (100 µm x 100 µm) that were not associated with any particles were counted as free-living bacteria, whereas bacteria that were on particles were counted as particle-associated. Sample filtration may bias counts due to free-living or particle-associated cells being hidden on the underside of particles, free-living bacteria settling on top of particles, or particle-associated cells dislodging. In the absence of any quantitative studies that have rigorously addressed this issue, we have assumed the net effect of these opposing methodological biases to be negligible in the present study.

### Heterotrophic bacterial production measurements

Community-wide heterotrophic bacterial production was measured using [_3_H] leucine incorporation into bacterial protein in the dark (Kirchman et al. 1985; Simon and Azam, 1989). At the end of the incubation with [_3_H]-leucine, cold trichloroacetic acid-extracted samples were filtered onto 3 µm filters that represented the leucine incorporation by particle-associated bacteria (>3.0 µm). Each filtrate was collected and filtered onto 0.2 µm filters and the activity therein represented incorporation of leucine by free-living bacteria (>0.2 µm-<3 µm). Measured leucine incorporation during the incubation was converted to bacterial carbon production rate using a standard theoretical conversion factor of 2.3 kg C per mole of leucine (Simon and Azam, 1989). Per-capita heterotrophic production was estimated by dividing heterotrophic production by the cell counts measured in each fraction.

### Preservation of bacterial filters in the field

Microbial biomass for the particle-associated (> 3 μm) and the free-living (3–0.22 μm) bacterial fraction was collected by sequential in-line filtration on 3 μm isopore polycarbonate (TSTP, 47 mm diameter, Millipore, Billerica, MA, USA) and 0.22 μm Express Plus polyethersulfone membrane filters (47 mm diameter, Millipore, MA, USA). We used 47 mm polycarbonate in-line filter holders (Pall Corporation, Ann Arbor, MI, USA) and an E/S portable peristaltic pump with an easy-load L/S pump head (Masterflex®, Cole Parmer Instrument Company, Vernon Hills, IL, USA). The total volume filtered varied from 0.8–2.2 L with a maximum filtration time of 16 minutes per sample. Filters were submerged in RNAlater (Ambion) in 2 mL cryovials, frozen in liquid nitrogen and transferred to a −80°C freezer until DNA extraction.

### DNA extraction, sequencing and processing

DNA extractions were performed using an optimized method based on the AllPrep DNA/RNA/miRNA Universal kit (Qiagen; McCarthy et al., 2015; details in supplementary methods). Extracted DNA was sequenced using Illumina MiSeq V2 chemistry 2 × 250 (500 cycles) of dual index-labelled primers that targeted the V4 hypervariable region of the 16S rRNA gene (515F/806R) (Caporaso et al., 2012; Kozich et al., 2013) at the Microbial Systems Laboratories at the University of Michigan Medical School in July 2016. RTA V1.17.28 and MCS V2.2.0 software were used to generate data. Fastq files were submitted to NCBI sequence read archive under BioProject accession number PRJNA412984. We analyzed the sequence data using MOTHUR V.1.38.0 (seed = 777; Schloss et al., 2009) based on the MiSeq standard operating procedure accessed on 3 November 2015 and modified with time (see data accessibility and supplemental methods). For classification of operational taxonomic units (OTUs), a combination of the Silva Database (release 123; Quast et al., 2013) and the freshwater TaxAss 16S rRNA database and pipeline (Rohwer et al., 2018, accessed August 18, 2016). All non-bacterial and chloroplast sequences were pruned out of the dataset and replicate samples were merged by summing sample sequencing read counts using the *merge_samples* function (phyloseq). A batch script for our protocol can be found in this project’s GitHub page in https://github.com/DenefLab/Diversity_Productivity/blob/master/data/mothur/mothur.batch.taxass.

### Estimating Diversity

To get the best estimate of each diversity metric, each sample was subsampled to 6,664 sequences (the smallest library size) with replacement and were averaged over 100 trials. Observed richness, Shannon entropy, and inverse Simpson’s index were calculated using the *diversity* function within the vegan (Oksanen et al., 2013) R package via the *estimate_richness* function in the phyloseq (McMurdie and Holmes, 2013) R package. Simpson’s Evenness was calculated by dividing the inverse Simpson’s index by the observed richness (Magurran, 2004). To calculate phylogenetic diversity, we first removed OTUs that had a count of 2 sequences or less throughout the entire dataset, as these are more prone to be artefacts originating from sequencing errors or the OTU clustering algorithm. Representative sequences of each of the 1,891 remaining OTUs were collected from the aligned fasta file produced within mothur, and header names in the mothur output fasta file were modified using bbmap (Bushnell, 2016) to only include the OTU name. A phylogenetic tree was created with FastTree using the GTR+CAT (general time reversible) model (Price et al., 2010). Mismatches between the species community data matrix and the phylogenetic tree were checked with the *match.phylo.comm* command (picante). Finally, both abundance-unweighted and -weighted phylogenetic diversity was estimated using specifications described in the next paragraph with the picante R package.

The most common phylogenetic diversity (PD) measure is Faith’s PD (Faith, 1992), however, this metric is very strongly correlated with species richness (**Figure S2**). Instead, the mean pairwise phylogenetic distance (or MPD) was calculated (*ses.mpd* function in the *Picante* R package (Kembell et al., 2010), null.model = “independentswap”). The MPD measures the average phylogenetic distance between all combinations of two taxa pulled from the observed community and compares it with a null community of equal richness pulled from the gamma diversity of all the samples (*see supplemental methods for more details*). Values higher than zero indicate phylogenetic evenness or overdispersion (higher phylogenetic diversity) while values less than zero indicate phylogenetic clustering (lower diversity) or that species are more closely related than expected according to the null community (Kembel, 2009). Thus, this phylogenetic metric is relative. From here, we will refer to the SESMPD as the “phylogenetic diversity” for simplicity and clarity.

### Statistical analysis

Further analysis of sequence data was performed in R version 3.4.2 (R Core Team 2017; *see supplemental methods for more details*). To test which variable(s) were the best predictors of community and per-capita heterotrophic production, we performed variable selection via a lasso regression (using the *glmnet* R package, alpha = 1, and lambda.1se as the tuning parameter (Friedman et al., 2010)) on all of the environmental, biodiversity, and principal component variables. To further validate the lasso regression results, we performed ordinary least squares (OLS) regressions on all variables, including the principal components (PCA) of the euclidean distances of the environmental data. We used the Akaike information criterion (AIC) (accessed with the *broom::glance()* command) to select the best performing OLS regression model.

### Data and code availability

Original fastq files can be found on the NCBI sequence read archive under BioProject accession number PRJNA412984. Processed data and code can be found on the GitHub page for this project at https://deneflab.github.io/Diversity_Productivity/ with the main analysis at https://deneflab.github.io/Diversity_Productivity/Final_Analysis.html

## Results

### Free-living communities had more cells but particle-associated communities had higher per-capita heterotrophic production

We observed an order of magnitude more cells per milliliter (p = 1 x 10-6, Figure 1A) and ∼2.5 times more community-wide heterotrophic production in the free-living fraction (p = 0.024, Figure 1B). However, when calculated per-capita, particle-associated bacteria were on average an order of magnitude more productive than free-living bacteria (p = 7 x 10-5, Figure 1C). Particle-associated and free-living cell abundances in samples taken from the same water sample did not correlate (Figure S3A). Heterotrophic production between corresponding free-living and particle-associated fractions from the same water sample were positively correlated for both community (Adjusted R_2_ = 0.40, p = 0.017; Figure S3B) and per-capita production rates (Adjusted R_2_ = 0.60, p = 0.003; Figure S3C).

**Figure 1:**
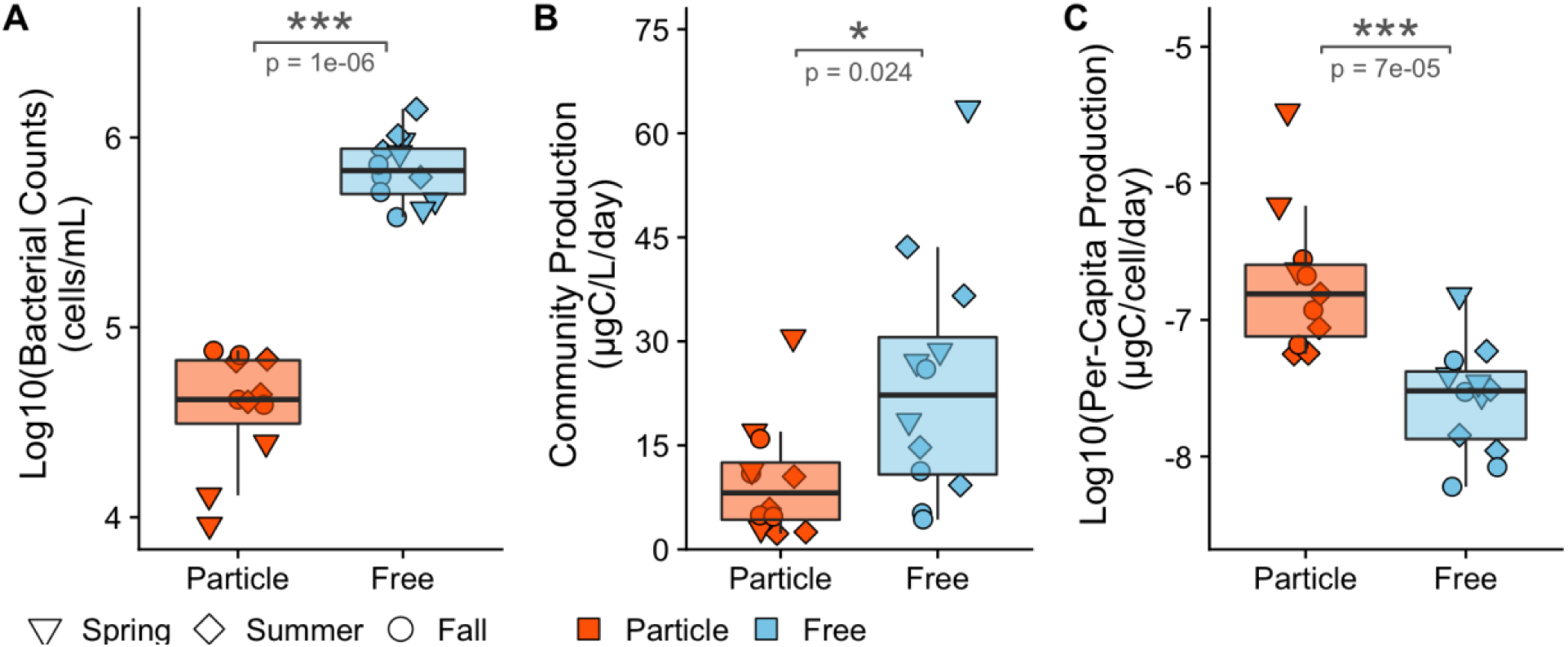
Bacterial counts, community-wide and per-capita heterotrophic production differ between microhabitats. Particle-associated and free-living samples were taken from four stations within Muskegon Lake during 2015 in May, July, and September. **(A)** Free-living bacteria were an order of magnitude (106 cells/mL) more abundant compared to particle-associated bacteria. **(B)** Free-living bacteria were more heterotrophically productive compared to particle-associated bacteria. **(C)** Particle-associated bacteria were disproportionately heterotrophically productive per cell compared to free-living bacteria.

### Particle-associated communities were more diverse in terms of observed richness and Shannon Entropy while free-living communities were more phylogenetically diverse

Across all samples, particle-associated bacterial communities were more diverse than free-living communities when considering richness and Shannon entropy (Figures 2A & S4A), but similar in the inverse Simpson’s index and Simpson’s evenness (Figure 2B & S4B).

**Figure 2:**
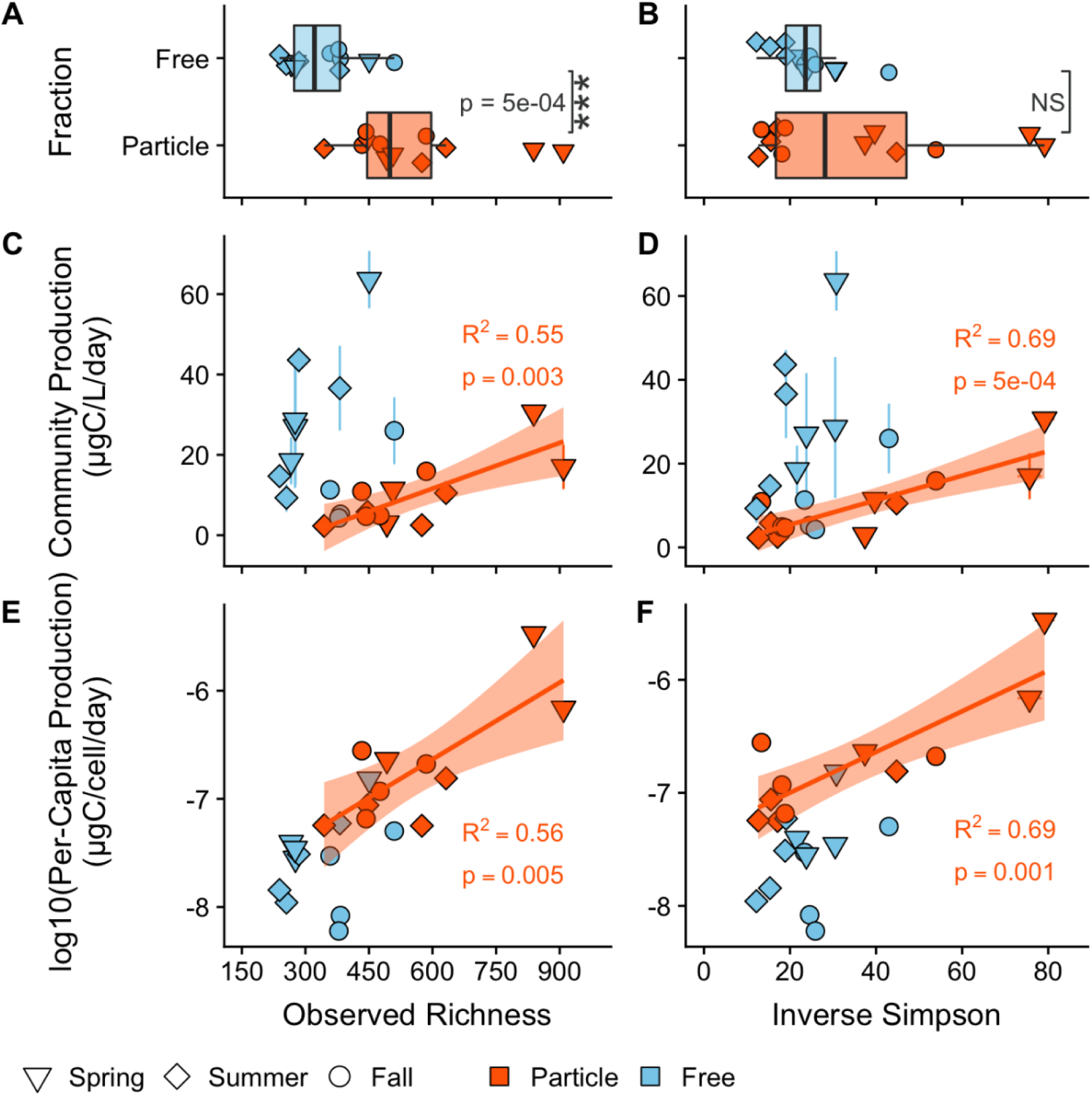
Richness and inverse Simpson correlate with heterotrophic productivity. Top panel: Differences in **(A)** the observed richness and **(B)** the inverse Simpson diversity metrics between particle-associated (orange) and free-living (blue) habitats. **Middle panel:** Biodiversity and community-wide heterotrophic production (ugC/L/day) relationships. The y-axis between **(C)** and **(D)** is the same, however, the x-axis represents **(C)** richness and **(D)** inverse Simpson. **Bottom panel:** Biodiversity and log10(per-capita heterotrophic production) (ugC/cell/day) relationships. The y-axis between **(E)** and **(F)** is the same, however, the x-axis represents **(E)** richness and **(F)** inverse Simpson’s index. Solid lines represent ordinary least squares models for the free-living (blue) and particle associated (orange) communities. All R_2_ values represent the adjusted R_2_ from an ordinary least squares model.

Particle-associated bacterial community richness was always higher than in free-living communities and was maintained across the four sampling stations in the lake (Figure S5A). Particle-associated samples at the river and Bear Lake stations were on average more OTU-rich than the outlet to Lake Michigan and the Deep stations. Additionally, the river station had almost twice the inverse Simpson’s value as compared with all other lake stations (Mean inverse Simpson Indices: Outlet = 23.6; Deep = 23.7; Bear = 35.3; River = 59.1; Figure S5A).

Particle-associated communities were more phylogenetically clustered than free-living communities based on unweighted phylogenetic diversity (p = 0.01, Figure 3A). Compared to other particle-associated samples, the outlet station that connects to oligotrophic Lake Michigan had a much larger unweighted phylogenetic diversity, indicating phylogenetic overdispersion (Figure S5A). Nevertheless, no sample across the entire dataset differed significantly from the null model with a significance threshold p-value of 0.05. There was no difference between weighted phylogenetic diversity in particle-associated versus free-living communities (Figure S5A).

**Figure 3:**
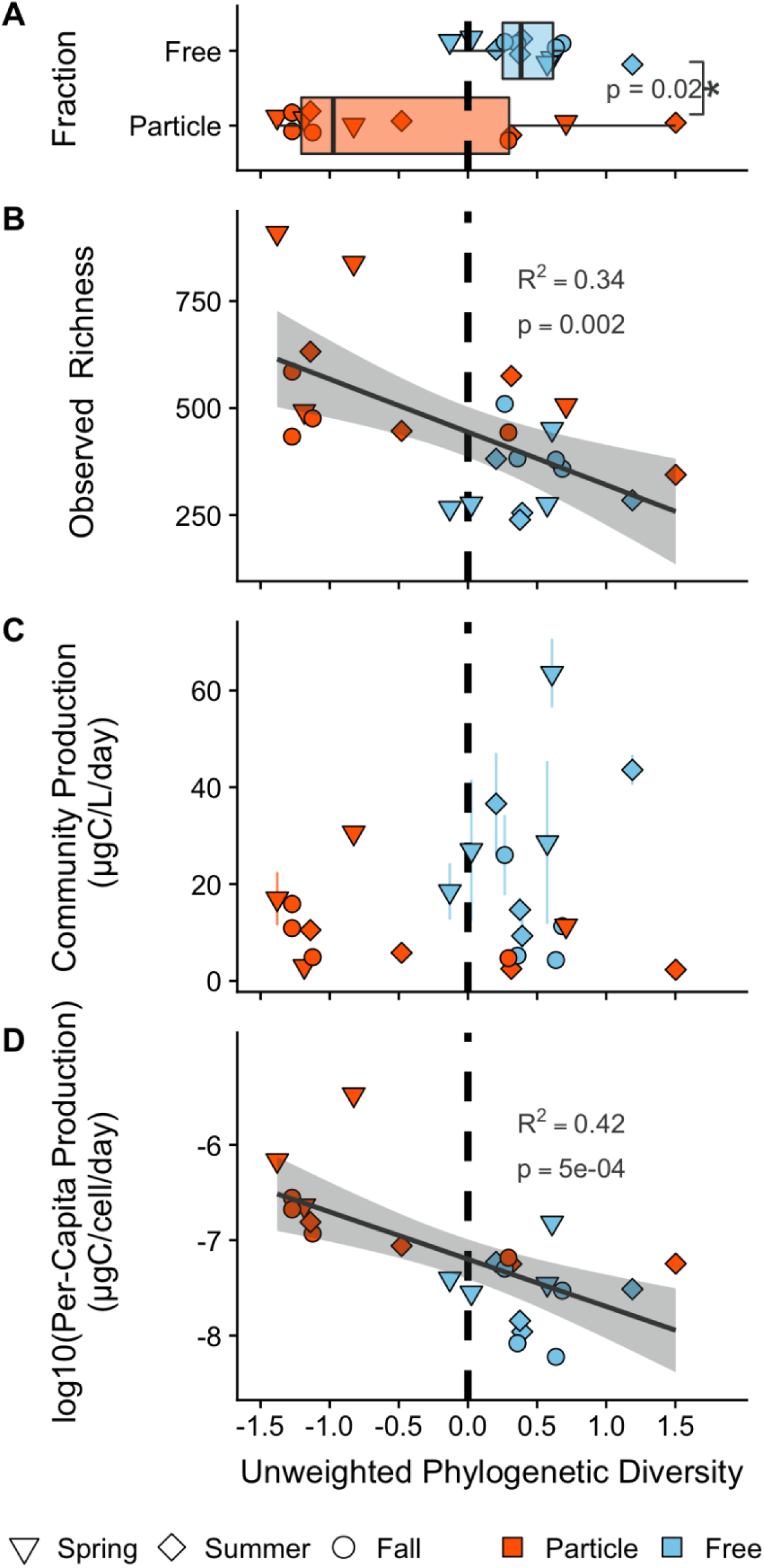
The relationship between heterotrophic productivity and unweighted phylogenetic diversity. (SESMPD; ses.mpd function in *picante* with null.model = “independentswap”). Positive phylogenetic diversity values represent communities that are phylogenetically diverse (*i.e.* overdispersed) while negative phylogenetic diversity values represent communities that are phylogenetically less diverse (*i.e.* clustered) compared to a null community with equal species richness. **(A)** Phylogenetic diversity was higher in free-living communities compared to particle-associated communities. **(B)** Negative relationship between observed richness and phylogenetic diversity. **(C)** Absence of phylogenetic diversity and community bulk heterotrophic production (µgC/L/day) relationships. **(D)** Negative phylogenetic diversity and per-capita heterotrophic production (µgC/cell/day) relationship. Linear models in figure **B** and **D** represent trends over all samples.

### Diversity-Productivity relationships were mostly observed in particle-associated communities

We analyzed BEF relationships for both community and per-capita production due to the distinct patterns of these two measures of heterotrophic production (Figure 1). There was a strong, positive, linear BEF relationship between community-wide (Figures 2C-D & S4C-D) and per-capita (Figures 2E-F & S4E-F) heterotrophic production and all richness and evenness diversity metrics in the particle-associated communities, while no BEF relationships were observed for the free-living communities. The inverse Simpson’s index explained the most amount of variation in community-wide heterotrophic production (Figure 2D; Adjusted R_2_ = 0.69, p = 5 x 10-4) and per-capita (Figure 2F; Adjusted R_2_ = 0.69, p = 0.001). When the two data points with the highest inverse Simpson’s index and heterotrophic production were removed from the regression (Figure 2D), the relationship was still significant and (Adjusted R_2_ = 0.37; p = 0.036), though not with richness (Adjusted R_2_ = 0.12; p = 0.17). These results are also robust across a range of minimum OTU abundance filtering thresholds (see *Sensitivity Analysis of Rare Taxa* in the supplemental methods and Figure S6) and hold up for all threshold levels in inverse Simpson and for richness until removal of OTUs observed 25 times (community-wide heterotrophic production) and 15 times (per-capita heterotrophic production).

When the particle-associated and free-living samples were combined together into one linear model to test an overall relationship between diversity and community-wide productivity, there was no relationship (richness: p = 0.86; Shannon: p = 0.99; inverse Simpson: p = 0.36), with the exception of a weak correlation for Simpson’s Evenness (Adjusted R_2_ = 0.12, p = 0.054). This further highlights the distinct BEF relationships across habitats. However, when particle-associated and free-living samples were combined together into one linear model to test an overall relationship between diversity and per-capita productivity, there was a strong relationship with observed richness (Adjusted R_2_ = 0.63, p = 3 x 10-6), which broke down as evenness was weighed more (Figure S7: Shannon: Adjusted R_2_ = 0.52, 6 x 10-5; Inverse Simpson: Adjusted R_2_ = 0.48, p = 2 x 10-4; Simpson’s Evenness: p = 0.48). Thus, the relationship between diversity and per-capita heterotrophic production was independent of habitat.

### Phylogenetic diversity correlated with per-capita heterotrophic production but not with community-wide production

Abundance-weighted phylogenetic diversity was not correlated with community or per-capita heterotrophic production (Figure S8C -S8D) and therefore no further analyses were performed with this diversity metric.

For unweighted phylogenetic diversity, we first determined the relationship between community richness and phylogenetic diversity. There was a moderate, negative, linear relationship when particle-associated and free-living samples were combined together into one linear model to test an overall relationship between unweighted phylogenetic diversity and observed richness (Figure 3B; Adjusted R_2_ = 0.35, p = 0.001). To further validate this trend, randomized communities were generated with an equal richness as the samples but with OTUs randomly picked across the dataset. The unweighted phylogenetic diversity was then calculated and regressed against each the randomized richness and there was no relationship (Figure S9; Adjusted R_2_ = -0.02, p = 0.44), verifying the negative relationship in the actual samples. When particle-associated and free-living samples were individually run in separate linear models to test for habitat-specific relationships between unweighted phylogenetic diversity and observed richness, no trend was found in either particle-associated or free-living models (Figure 3B; Particle: Adjusted R_2_ = 0.14, p = 0.12; Free = Adjusted R_2_ = -0.10, p = 0.97).

We then assessed the relationship between phylogenetic diversity and heterotrophic productivity. Particle-associated and free-living phylogenetic diversities did not have individual effects on community-wide or per-capita heterotrophic production. When data from both habitats were combined into a single regression, there was no correlation between phylogenetic diversity and community-wide heterotrophic production (Figure 3C). However, a negative correlation was found when particle-associated and free-living samples were combined into one linear model to test an overall relationship between unweighted phylogenetic diversity and per-capita heterotrophic production (Figure 3D; Adjusted R_2_ = 0.42, p = 5 x 10-4). Therefore, these two results in combination indicated that communities composed of more phylogenetically similar OTUs had a higher per-capita heterotrophic production rate.

### Diversity, and not environmental variation, was the best predictor of particle-associated heterotrophic production

To identify variables that best predicted community-wide and per-capita heterotrophic production (*i.e.* remove variables that were correlated with each other and/or uninformative variables), we performed lasso regression with all samples and individually with particle-associated and free-living samples. For prediction of community-wide heterotrophic production, only the inverse Simpson’s index was selected for particle-associated samples whereas pH and PC5 were selected for free-living samples, and no variables were selected when all samples were included in the lasso regression. In contrast, for per-capita heterotrophic production, temperature and the inverse Simpson’s index were selected for particle-associated samples whereas pH was the only predictor for free-living samples, and observed richness was the only predictor for all samples (plotted in Figure S7A). Therefore, the best model for particle-associated microhabitats *always* included inverse the inverse Simpson’s index whereas free-living samples only included environmental variables, such as pH.

To further verify that there were no confounding impacts of seasonal and environmental variables on community-wide and per-capita heterotrophic production, we performed ordinary least square (OLS) regressions and a dimension-reduction analysis of the environmental variables through a principal components analysis (Table S1 & S2; Figure S10). Specifically, the first 2 environmental axes explained ∼70% of the environmental variation in the sampling sites (Figure S10). Next, we predicted community-wide and per-capita heterotrophic production with all environmental variables and the first six principal components as predictor variables with individual particle-associated and free-living samples, and combined (*i.e.* all samples) models (Table S1 & S2). The best single predictor of community-wide heterotrophic production was inverse Simpson for particle-associated samples (AIC = 74.34; Adjusted R_2_ = 0.69), pH for the free-living samples (AIC =98.43; Adjusted R_2_ = 0.49, p = 0.006), and pH for all samples (AIC = 192.16; Adjusted R_2_ = 0.35) (Table S1). Whereas, the best single predictor of per-capita heterotrophic production was inverse Simpson for particle-associated samples (AIC = 8.29; Adjusted R_2_ = 0.69), pH for the free-living samples (AIC = -2.39; Adjusted R_2_ = 0.78), and observed richness for all samples (AIC = 24.72; Adjusted R_2_ = 0.63) (Table S2). Thus, the OLS regressions are in agreement with the lasso regressions.

## Discussion

We examined bacterial biodiversity-ecosystem function (BEF) relationships in relation to two microhabitats within freshwater lakes: particulate matter and the surrounding water. First, we found that community-wide and per-capita heterotrophic productivity of particle-associated but not free-living bacterial communities showed a positive, linear BEF relationship with both richness and evenness contributing. Second, particle-associated heterotrophic production was better explained by diversity (*i.e.* inverse Simpson’s index) than by environmental parameters.

Third, across both particle-associated and free-living communities, higher richness was associated with lower phylogenetic diversity which, in turn, was associated with higher per-capita heterotrophic bacterial production but not associated with community-wide heterotrophic production.

Microbes have a large diversity of metabolisms and the choice of which to focus on may inherently affect the BEF relationship. Indeed, “narrow” metabolic processes that are catalyzed by a small subset of taxa within bacterial communities, such as some nitrogen and sulfur cycling, have been found to display BEF relationships (Levine et al., 2011; Delgado-Baquerizo et al., 2016). In contrast, for “broad” processes that are performed by the majority of taxa within a bacterial community, such as heterotrophic production (*i.e.* focus of the present study) and respiration, functional redundancy appears to weaken or remove the presence of BEF relationships (Griffiths et al., 2000; Langenheder et al., 2006; Wertz et al., 2006; Levine et al., 2011; Peter et al., 2011, Galand et al, 2015). These findings are in line with the absence of a BEF relationship for free-living bacterial communities in our study.

However, the above results and hypotheses surrounding narrow and broad processes are in conflict with the strong BEF relationship we observed in particle-associated bacterial communities. As such, our study signifies that microhabitats or habitat heterogeneity can influence bacterial BEF relationships, in agreement with previous research in eukaryotic systems across a variety of ecosystems (Tylianakis et al., 2008; Cardinale 2011; Zeppilli et al., 2016). A study using controlled stream mesocosms by Cardinale (2011) found that niche complementarity effects are particularly important in more heterogeneous environments. In more heterogeneous streams, algal populations used different nutrients and avoided direct competition for resources, resulting in unique species occupying distinct and local microhabitats (Cardinale 2011). An experimental study by Gravel et al. (2011) showed that BEFs depend on the legacy of previous evolutionary events. Specifically, they found that after several hundred generations of evolution on a variety of carbon substrates, generalist bacteria were more productive because of their more efficient exploitation of the environmental heterogeneity. Finally, a recent study on freshwater lake bacterial communities found a positive correlation between OTU evenness and the number of dissolved organic matter (DOM) components (though the study did not evaluate ecosystem function), suggesting that DOM resource heterogeneity may increase the diversity of bacterial communities by creating equity among bacterial species (Muscarella et al., 2019).

Our observational study could not directly test the role of niche complementarity effects. However, support for niche complementarity alone or in combination with species selection as the mechanism underlying the BEF relationship in particle-associated habitats is provided by the inverse Simpson’s index being the strongest predictor of community-wide heterotrophic production. As the inverse Simpson’s index represents a measure of species dominance, it is strongly affected by the evenness of abundant species. Communities that are more even have an increased likelihood for complementary species to neighbor each other.

In our study, there are several reasons why heterogeneity of particulate matter may allow for niche complementarity effects to occur and result in BEF relationships. First, particles have a two-fold layer of heterogeneity as they (A) may be composed of different substrates such as organic matter from terrestrial or aquatic environments and either heterotrophically or photosynthetically derived (Grossart, 2010), and (B) each particle may comprise physicochemical gradients as well (Simon et al., 2002). Second, microbial interactions are more likely to occur between cells aggregated on particles as the interaction distances are usually much shorter (Cordero & Datta, 2016) compared to free-living bacterial cells. In fact, genes mediating social interactions, such as motility, adhesion, cell-to-cell transfer, antibiotic resistance, mobile element activity, and transposases, have been found to be more abundant in marine particles than compared to the surrounding water (Ganesh et al., 2014). Third, a recent study by Ebrahimi et al. (bioRxiv) found that dense patches of bacterial cells on model marine chitin particles promoted cross-feeding of oligosaccharides when particles were recalcitrant.

The importance of niche complementarity in microbial communities can also be deduced from recent findings in the field of microbiology, which have shown widespread metabolic interdependence among bacterial community members. First, a 2016 study that reconstructed 2,540 draft genomes of microbes found that most bacteria specialize in one particular step in sulfur and nitrogen pathways and “hand-off” their metabolic byproducts to nearby organisms (Anantharaman et al., 2016). It is likely that metabolic hand-offs, a specific form of bacterial facilitation, will occur more in particle-associated compared to free-living communities. Indeed, Datta et al.’s (2016) work on model marine chitin particles found that taxa that are incapable of breaking down particles and instead rely on carbon produced by primary degraders thrive in later phases of particle degradation, a repeatable result for three other polysaccharide substrates (Enke et al., 2019). Second, Lilja and Johnson (2016) demonstrated that different microbial cell types eliminate inter-enzyme competition by cross feeding, which increases substrate consumption by allowing intracellular resources to go towards a single enzyme, rather than having two enzymes that perform two separate reactions compete for nutrients within a cell. Third, some bacteria are unable to grow in laboratory cultures unless they are in co-culture with other organisms, which may be due to metabolic hand-offs or growth factors such as siderophores or catalases (Stewart, 2012).

Considering that (i) closely related taxa share more genes and metabolic pathways than distantly related bacterial taxa (Konstantinidis & Tiedje, 2005; Kim et al., 2014) and (ii) bacteria commonly have incomplete metabolic pathways, it may be possible that closely related bacteria are most likely to exchange their metabolic byproducts. This may be why we found that new taxa added to the community represented taxonomic clades similar to or already present in the community, and that these communities with lower phylogenetic diversity (relative to expected) had higher productivities. This result agrees with a recent study using freshwater algae and vascular plants that found that co-cultures of more similar species were more productive (Narwani et al., 2017). Similarly, a study of bacterial communities inhabiting Mediterranean soils showed that plots containing more recently diverged lineages had higher ecosystem function levels than when more distantly related lineages were present (Goberna & Verdú, 2018). However, other bacteria-focused studies found higher levels of antagonism with more closely related taxa (Russel et al., 2017) and more bacterial productivity (measured through colony forming units per mL) with more distantly related taxa (Venail and Vives, 2013). Though, both of these studies were performed in the lab with r-selected (*i.e.* copiotrophic) species grown in stable, warm, aerobic, agar plate conditions. Thus, Venail and Vives (2013) and Russel et al. (2017) inherently break up potential interdependent relationship between bacteria either by creating artificial communities or evaluating pairwise interactions and remove the natural effect of spatial heterogeneity, environmental fluctuations, and the rest of the bacterial community.

Previous studies on bacterial BEF relationships have used three approaches to manipulate bacterial diversity (Krause et al., 2014): (1) removal of taxa (*e.g.* dilution to extinction) in which complex communities are simplified (Franklin et al, 2001; Wertz et al., 2006; Peter et al., 2011; Philippot et al., 2013; see Roger et al., 2016 for a review of this approach; fragmentation or knockout in Bell, 2019), (2) addition of taxa (*e.g.* manually assembled communities) in culture (Tan et al., 2012; Salles et al., 2009; invasion or coalescence in Bell, 2019), or (3) natural or manipulated environmental communities (Griffiths et al., 2000; Levine et al., 2011; Galand et al., 2015; Rivett & Bell, 2018). We took the third approach in this study. In contrast to the other two approaches, this had the benefit of (a) maintaining high diversity with both abundant and rare taxa, (b) including both r- and k-selected organisms, (c) allowing natural environmental and ecological forcings to shape the community, and (d) evaluating BEF relationships in diversity and productivity ranges that reflect natural communities. Admittedly, three inherent weaknesses to our approach were that (a) we cannot measure all the potential variables that influence heterotrophic productivity, which are especially and inherently difficult to measure within particles, (b) we only have 24 samples for a 12 versus 12 study, and (c) our analysis is correlational and we cannot manipulate the system to unequivocally separate causes and consequences of bacterial production. For example, strong correlations with heterotrophic production and pH in the free-living samples (Table S1 & S2) may point to pH being a consequence of rather than a cause of varying production levels. This is because bacterial production and bacterial respiration are positively correlated (del Giorgio & Cole, 1998) and with increased respiration, pH may decrease due to CO_2_ dissolution into the water.

Finally, we acknowledge that the typical sampling of bacterial communities and analysis using DNA sequencing reflects *all* bacteria present in the community and not necessarily only the *active* members of the community contributing to a given ecosystem function. In freshwater systems, up to 40% of cells from the total community have been found to be inactive or dormant (Jones and Lennon, 2010). In addition, leucine incorporation is not universal across all taxa (Salcher et al, 2013). In this context, it is interesting to reflect on the richness in absence of function (i.e. x-intercept) of the observed BEF relationship which is 295 (Figure 2C). This could be interpreted as a baseline level of 295 particle-associated OTUs that are inactive (either dead or dormant cells or environmental DNA) or incapable of incorporating leucine. This value represents 35-85% of the total particle-associated communities and may obscure the *actual* diversity (and BEF relationship) of the bacterial community (Carini et al., 2016).

In conclusion, we show that increased bacterial diversity leads to increased bacterial heterotrophic production in particle-associated but not in free-living communities. As such, we extend the validity of principles of the impact of microhabitat on BEF relationships from Eukarya to Bacteria, contributing to current efforts to integrate ecological theories into the field of microbiology (Barberán et al., 2014). Additionally, we show that communities with low phylogenetic diversity have higher per-capita heterotrophic production rates, which we hypothesize to be related to genome evolutionary patterns specific to bacteria that result in the dependence on metabolic hand-offs. The unique nature of BEF relationships across particle-associated and free-living habitats agrees with the distinct community assembly and functional partitioning previously described between these two aquatic habitats (Bižić-Ionescu et al., 2014; Ganesh et al., 2014; Mohit et al., 2014; Schmidt et al., 2016; Balmonte et al., 2018). Future studies should focus on going beyond observations to predicting changes in ecosystem function based on community changes (Bell, 2019).

## Acknowledgements

This work was supported by the National Science Foundation Graduate Research Fellowship Grant No. DGE 1256260 (MLS), the University of Michigan Office for Research MCubed program (VJD), the American Society of Microbiology-Undergraduate Research Fellowship, the University of Michigan Honors Summer Fellowship, and the Beckman Scholars Program (EC). RP was supported by Ghent University (BOFDOC2015000601) and a Sofina Gustave-Boël grant from the Belgian American Educational Foundation. We are grateful to the crew of the R/V W.G. Jackson and the Grand Valley State University Robert B. Annis Water Resources Institute science staff, and the generous help we received in the field from Amelia Waters and Daniel S.W. Katz. Thank you to Kyle Buffin and Amadeus Twu for help with DNA extractions. Finally, we thank Deborah Goldberg, George Kling, and members of the Denef, Dick, and Duhaime laboratories for their comments on the manuscript.

## References

1. Anantharaman, K. et al. 2016. Thousands of microbial genomes shed light on interconnected biogeochemical processes in an aquifer system. Nature Communications 7:13219.

2. Balmonte, J.P., Teske, J.P., and C. Arnosti. 2018. Structure and function of high Arctic pelagic, particle-associated and benthic bacterial communities. 20(8):2941–2954.

3. Barberán, A., E. O. Casamayor, and N. Fierer. 2014. The microbial contribution to macroecology. Frontiers in Microbiology 5:1–8.

4. Bell, T. 2019. Next-generation experiments linking community structure and ecosystem functioning. Environmental Microbiology Reports, 11(1):20–22.

5. Bižić-Ionescu, B., Zeder, M., Ionescu. D., Orlic, S., Fuchs, B.M., Grossart, H.P., and R. Amann. 2014. Comparison of bacterial communities on limnic versus coastal marine particles reveals profund differences in colonization. Environmental Microbiology doi:10.1111/1462-2920.12466.

6. Blaser, M. J. 2014. Missing Microbes: How the Overuse of Antibiotics Is Fueling Our Modern Plagues. Henry Holt and Company LLC, New York 35:261.

7. Bushnell B. 2016. BBMap short read aligner. https://sourceforge.net/projects/bbmap/.

8. Cadotte, M. W., B. J. Cardinale, and T. H. Oakley. 2008. Evolutionary history and the effect of biodiversity on plant productivity. Proceedings of the National Academy of Sciences 105:17012–17017.

9. Caporaso, J. G. et al. 2012. Ultra-high-throughput microbial community analysis on the Illumina HiSeq and MiSeq platforms. The ISME Journal 6:1621–1624.

10. Cardinale, B. J. 2011. Biodiversity improves water quality through niche partitioning. Nature 472:86–89.

11. Cardinale, B. J. et al. 2012. Biodiversity loss and its impact on humanity. Nature 489:326–326.

12. Carini, P., P. J. Marsden, J. W. Leff, E. E. Morgan, M. S. Strickland, and N. Fierer. 2016. Relic DNA is abundant in soil and obscures estimates of soil microbial diversity. Nature Microbiology 2:16242.

13. Cordero, O. X., and M. S. Datta. 2016. ScienceDirect Microbial interactions and community assembly at microscales. Current Opinion in Microbiology 31:227–234.

14. Cotner, J. B., and B. A. Biddanda. 2002. Small players, large role: Microbial influence on biogeochemical processes in pelagic aquatic ecosystems. Ecosystems 5:105–121.

15. Crump, B.C., Armbrust, E.V., and J.A. Baross. 1999. Phylogenetic analysis of particle-attached and free-living bacterial communities in the Columbia River, its estuary, and the adjacent coastal ocean. Applied and Environmental Microbiology 65(7):3192–3204.

16. Datta, M. S., Sliwerska, E., Gore, J., Polz, M. F. and O. X. Cordero. 2016. Microbial interactions lead to rapid micro-scale successions on model marine particles. Nature Communications 7:11965.

17. Delgado-Baquerizo, M., L., et al. 2016. Lack of functional redundancy in the relationship between microbial diversity and ecosystem functioning. Journal of Ecology 104:936–946.

18. del Giorgio, P. A., and J. J. Cole. 1998. Bacterial Growth Efficiency in Natural Aquatic Systems. Annual Review of Ecology and Systematics 29:503–541.

19. Duffy, J.E., Godwin, C.M., and B.J. Cardinale. 2017. Biodiversity effects in the wild are common and as strong as key drivers of productivity. Nature 549:261–264.

20. Ebrahimi, A., Schwartzman, J., and O.X. Cordero. Cooperation and spatial self-organization determine ecosystem function for polysaccharide-degrading bacteria. bioRxiv:640961.

21. Enke, T.N., et al. 2019. Modular assembly of polysaccharide-degrading marine microbial communities. Current Biology 29:1–8.

22. EPA. 1993. Methods for the determination of inorganic substances in environmental samples. USEPA 600/R-93/100.

23. Evans, R., Alessi, A.M., Bird, S., McQueen-Mason, S.J., Bruce, N.C., and M.A. Brockhurst. 2017. Defining the functional traits that drive bacterial decomposer community productivity. The ISME Journal 11:1680–1687.

24. Faith, D. P. 1992. Conservation evaluation and phylogenetic diversity. Biological Conservation 61:1–10.

25. Ferrier S, K. N. Ninan, P. Leadley, R. Alkemade GK, M. Moraes R., & E. Y. Mohammed and Y. Trisurat. 2016. Overview and vision. In IPBES 2016: The methodological assessment report on scenarios and models of biodiversity and ecosystem services. S. Ferrier, K. N. Ninan, P. Leadley, R. Alkemade, L. A. Acosta, H. R. Akçakaya, L. Brotons, W. W. L. Cheung, V. Christensen, K. A. Harhash, J. KabuboMariara, C. Lundquist, M. Obersteiner, H. M. Pereira, G. Peterson, R. Pichs-Madruga, N. Ravindranath, C. Rondinini and B.A. Wintle (eds.), Bonn, Germany, Secretariat of the Intergovernmental Science-Policy Platform for Biodiversity and Ecosystem Services.

26. Flynn, D.F.B., Mirotchnick, N., Jain, M., Palmer, M.I., and S. Naeem. 2011. Functional and phylogenetic diversity as predictors of biodiversity–ecosystem-function relationships. Ecology 92(8): 1573–1581.

27. Franklin, R.B., Garland, J.L., Bolster, C.H., and A.L. Mills. 2001. Impact of dilution on microbial community structure and functional potential: Comparison of numerical simulations and batch culture experiments. Applied and Environmental Microbiology 67(2):702–712.

28. Friedman, J., T. Hastie, and R. Tibshirani. 2010. Regularization Paths for Generalized Linear Models via Coordinate Descent. Journal of Statistical Software 33:1–22.

29. Fritschie, K. J., B. J. Cardinale, M. A. Alexandrou, and T. H. Oakley. 2014. Evolutionary history and the strength of species interactions: Testing the phylogenetic limiting similarity hypothesis. Ecology 95:1407–1417.

30. Galand, P. E., I. Salter, and D. Kalenitchenko. 2015. Ecosystem productivity is associated with bacterial phylogenetic distance in surface marine waters. Molecular Ecology 24:5785–5795.

31. Ganesh, S., D. J. Parris, E. F. DeLong, and F. J. Stewart. 2014. Metagenomic analysis of size-fractionated picoplankton in a marine oxygen minimum zone. The ISME journal 8:187–211.

32. Goberna, M. and M. Verdú. 2018. Phylogenetic-scale disparities in the soil microbial diversity–ecosystem functioning relationship. The ISME Journal 12:2152–2162.

33. Gravel, D., Bell, T., Barbera, C., Bouvier, T., Pommier, T., Venail, P., and N. Mouquet. 2011. Experimental niche evolution alters the strength of the diversity-productivity relationship. Nature 469:89–92.

34. Griffiths, B. S. et al. 2000. Ecosystem response of pasture soil communities to fumigation-induced microbial diversity reductions: an examination of the biodiversity-ecosystem function relationship. Oikos 90:279–294.

35. Grossart, H.P., Tang, K.W., Kiørboe, T., and H. Ploug. 2007. Comparison of cell-specific activities between free-living and attached bacteria using isolates and natural assemblages. FEMS Microbiology Letters 266:194–200.

36. Grossart, H. P. 2010. Ecological consequences of bacterioplankton lifestyles: Changes in concepts are needed. Environmental Microbiology Reports 2:706–714.

37. Hobbie, J. E., Daley, R. J., and S. Jasper. 1977. Use of nuclepore filter counting bacteria by fluorescence microscopy. Applied and Environmental Microbiology 33:1225–1228.

38. Jiang, L., J. Tan, and Z. Pu. 2010. An Experimental Test of Darwin’s Naturalization Hypothesis. The American Naturalist 175:415–423.

39. Jones, S. E., and J. T. Lennon. 2010. Dormancy contributes to the maintenance of microbial diversity. Proceedings of the National Academy of Sciences 107:5881–5886.

40. Kembel, S. W. 2009. Disentangling niche and neutral influences on community assembly: Assessing the performance of community phylogenetic structure tests. Ecology Letters 12:949–960.

41. Kembel, S.W. et al. 2010. Picante: R tools for integrating phylogenies and ecology. Bioinformatics 26:1463–1464.

42. Kim, M., Oh, H. S., Park, S. C., and J. Chun. 2014. Towards a taxonomic coherence between average nucleotide identity and 16S rRNA gene sequence similarity for species demarcation of prokaryotes. International Journal of Systematic and Evolutionary Microbiology 64:346–351.

43. Kirchman, D., E. K’nees, and R. Hodson. 1985. Leucine incorporation and its potential as a measure of protein synthesis by bacteria in natural aquatic systems. Applied and Environmental Microbiology 49:599–607.

44. Kirwan, L. et al. 2007. Evenness drives consistent diversity effects in intensive grassland systems across 28 European sites. Journal of Ecology 95:530–539.

45. Konstantinidis, K. T. and J. M. Tiedje. 2005. Genomic insights that advance the species definition for prokaryotes. Proceedings of the National Academy of Sciences 102:2567–2572.

46. Kozich, J. J., S. L. Westcott, N. T. Baxter, S. K. Highlander, and P. D. Schloss. 2013. Development of a dual-index sequencing strategy and curation pipeline for analyzing amplicon sequence data on the miseq illumina sequencing platform. Applied and Environmental Microbiology 79:5112–5120.

47. Krause, S. et al. 2014. Trait-based approaches for understanding microbial biodiversity and ecosystem functioning. Frontiers in Microbiology 5:1–10.

48. Langenheder, S., E. S. Lindström, and L. J. Tranvik. 2006. Structure and Function of Bacterial Communities Emerging from Different Sources under Identical Conditions. Applied and Environmental Microbiology 72:212–220.

49. Levine, U. Y., T. K. Teal, G. P. Robertson, and T. M. Schmidt. 2011. Agriculture’s impact on microbial diversity and associated fluxes of carbon dioxide and methane. The ISME Journal 5:1683–1691.

50. Lilja, E. E. and D. R. Johnson. 2016. Segregating metabolic processes into different microbial cells accelerates the consumption of inhibitory substrates. The ISME Journal 10:1–11.

51. Magurran, A. E. 2004. Chapter four: An index of diversity in Measuring Biological Diversity, Wiley-Blackwell, Hoboken, NJ.

52. McMurdie, P. J., and S. Holmes. 2013. phyloseq: An R Package for Reproducible Interactive Analysis and Graphics of Microbiome Census Data. PLoS ONE 8:e61217.

53. Mohit, V., Archambault, P., Toupoint, N., and C. Lovejoy. 2014. Phylogenetic differences in attached and free-living bacterial communities in a temperate coastal lagoon during summer, revealed via high-throughput 16S rRNA gene sequencing. Applied and Environmental Microbiology 80: 2071–2083.

54. Muscarella, M.E., Boot, C. M., Broeckling, C. D., and J. T. Lennon. 2019. Resource heterogeneity structures aquatic bacterial communities. The ISME Journal 13:2183–2195.

55. Narwani, A. et al. 2017. Ecological interactions and coexistence are predicted by gene expression similarity in freshwater green algae. Journal of Ecology 105:580–591.

56. Oksanen, A. J. et al. 2015. vegan: Community Ecology Package. R package version 2.3–0.

57. Peter, H., S. Beier, S. Bertilsson, E. S. Lindström, S. Langenheder, and L. J. Tranvik. 2011. Function-specific response to depletion of microbial diversity. The ISME Journal 5:351–361.

58. Philippot, L. et al. 2013. Loss in microbial diversity affects nitrogen cycling in soil. The ISME Journal 7:1609–1619.

59. Price, M. N., P. S. Dehal, and A. P. Arkin. 2010. FastTree 2 – Approximately maximum-likelihood trees for large alignments. PLoS ONE 5.

60. Quast, C. et al. 2013. The SILVA ribosomal RNA gene database project: Improved data processing and web-based tools. Nucleic Acids Research 41:590–596.

61. R Core Team. 2017. R: A Language and Environment for Statistical computing. Vienna, Austria: R Foundation for Statistical Computing. https://www.R-project.org/.

62. Reich, P.B. et al. 2004. Species and functional group diversity independently influence biomass accumulation and its response to CO_2_ and N. Proceedings of the National Academy of Sciences 101(27):10101–10106.

63. Rivett, D.W. and T. Bell. 2018. Abundance determines the functional role of bacterial phylotypes in complex communities. Nature Microbiology:s41564–018-0180-0.

64. Roger, F., S. Bertilsson, S. Langenheder, O. A. Osman, and L. Gamfeldt. 2016. Effects of multiple dimensions of bacterial diversity on functioning, stability and multifunctionality. Ecology 97:2716–2728.

65. Rohwer, R. R., J. J. Hamilton, R. J. Newton, and K. D. McMahon. 2018. TaxAss: Leveraging a custom freshwater database achieves fine-scale taxonomic resolution. mSphere 3(5):e00327–18.

66. Russel, J., H. L. Røder, J. S. Madsen, M. Burmølle, and S. J. Sørensen. 2017. Antagonism correlates with metabolic similarity in diverse bacteria. Proceedings of the National Academy of Sciences:201706016.

67. Salcher, M.M., Posch, T., and J. Pernthaler. 2013. *In situ* substrate preferences of abundant bacterioplankton populations in a prealpine freshwater lake. The ISME Journal 7:896–907.

68. Salles, J. F. et al. 2009. Community niche predicts the functioning of denitrifying bacterial assemblages. Ecology, 90:3324–3332.

69. Schloss, P. D. et al. 2009. Introducing mothur: Open-source, platform-independent, community-supported software for describing and comparing microbial communities. Applied and Environmental Microbiology 75:7537–7541.

70. Schmidt, M.L., J.D. White, and V. J. Denef. 2016. Phylogenetic conservation of freshwater lake habitat preference varies between abundant bacterioplankton phyla. Environmental Microbiology 18(4):1212–1226.

71. Shade, A., S. E. Jones, and K. D. McMahon. 2008. The influence of habitat heterogeneity on freshwater bacterial community composition and dynamics. Environmental Microbiology 10:1057–1067.

72. Simon, M., and F. Azam. 1989. Protein content and protein synthesis rates of planktonic marine bacteria. Marine Ecology Progress Series 51:201–213.

73. Simon, M., H. P. Grossart, B. Schweitzer, and H. Ploug. 2002. Microbial ecology of organic aggregates in aquatic ecosystems. Aquatic Microbial Ecology 28:175–211.

74. Singh, B. K. et al. 2014. Loss of microbial diversity in soils is coincident with reductions in some specialized functions. Environmental Microbiology 16:2408–2420.

75. Steinman, A. D., Ogdahl, M., Rediske, R., Ruetz III, C.R., Biddanda, B. A., and L. Nemeth. 2008. Current status and trends in Muskegon Lake, Michigan. Journal of Great Lakes Research 34:169–188.

76. Stewart, E. J. 2012. Growing unculturable bacteria. Journal of Bacteriology 194:4151–4160.

77. Tan, J., Z. Pu, W. A. Ryberg, and L. Jiang. 2012. Species phylogenetic relatedness, priority effects, and ecosystem functioning. Ecology 93:1164–1172.

78. Thomas, C. D. et al. 2004. Extinction risk from climate change. Nature 427:145–8.

79. Tilman, D., F. Isbell, and J. M. Cowles. 2014. Biodiversity and Ecosystem Functioning. Annual Review of Ecology, Evolution, and Systematics 45:471–493.

80. Tylianakis, J. M. et al. 2008. Resource heterogeneity moderates the biodiversity-function relationship in real world ecosystems. PLoS Biology 6:0947–0956.

81. Venail, P. A., A. Narwani, K. Fritschie, M. A. Alexandrou, T. H. Oakley, and B. J. Cardinale. 2014. The influence of phylogenetic relatedness on species interactions among freshwater green algae in a mesocosm experiment. Journal of Ecology 102:1288–1299.

82. Venail, P. A., and M. J. Vives. 2013. Phylogenetic distance and species richness interactively affect the productivity of bacterial communities. Ecology 94:2529–2536.

83. Wake, D. B., and V. T. Vredenburg. 2008. Are we in the midst of the sixth mass extinction? A view from the world of amphibians. Proceedings of the National Academy of Sciences 105:11466–11473.

84. Wertz, S., V., et al. 2006. Maintenance of soil functioning following erosion of microbial diversity. Environmental Microbiology 8:2162–2169.

85. Wilsey, B.J. and H. W. Polley. 2004. Realistically low species evenness does not alter grassland species richness productivity relationships. Ecology 85(10):2693–2700.

86. Wilsey, B.J. and C. Potvin. 2000. Biodiversity and ecosystem functioning: Importance of species evenness in an old field. Ecology 81(4):887–892.

87. Wittebolle, L. et al. 2009. Initial community evenness favors functionality under selective stress. Nature 458(2)623–626.

88. Zeppilli, D., A. Pusceddu, F. Trincardi, and R. Danovaro. 2016. Seafloor heterogeneity influences the biodiversity–ecosystem functioning relationships in the deep sea. Scientific Reports 6:26352.

89. Zhou, J., S., et al. 2008. Spatial scaling of functional gene diversity across various microbial taxa. Proceedings of the National Academy of Sciences 105:7768–7773.

