## Supplemental Information for "Microhabitats shape diversity-productivity relationships in freshwater bacterial communities"

### Supplemental Methods

#### *Map of Muskegon Lake*

The Muskegon Lake map (Figure S1) was created by using the pre-existing 2006 National Hydrography Dataset (NHD) GIS feature data for the Muskegon Lake shoreline was adjusted to an updated 2008 format using ESRI™ ArcGIS software and high resolution (6” pixel) leaf-off aerial orthophotography. Historic mean water level for Lake Michigan was then used to estimate Muskegon Lake water level for April 2008 at 176.4 meters. This water level was used as the base elevation and adjustment point for correcting all relevant lake bathymetric data, taken from the recently, published February 2008 NOAA electronic bathymetric chart for Muskegon Lake. The corrected GIS shoreline boundary and the supplemental NOAA bathymetric point data were used to generate a new bathymetric grid (raster) feature for Muskegon Lake from which, contours were created at 2m depth intervals. This bathymetric map was then laid under a Google Earth image of Muskegon Lake and the immediately surrounding area.

#### *DNA Extraction*

In summary, the filters were first washed with phosphate-buffered saline (pH 7.4) and folded (cell-side in) to minimize cell loss and to remove RNAlater, which inhibits DNA yields. Then the filters were placed in a 2 mL tube with 600 µL of buffer RLT plus (Qiagen) and incubated for 90 min at room temperature using a vortex on medium setting (5 of 10). After incubation, tubes were vortexed on high for 10 min. The lysate was transferred to a QiaShredder column (Qiagen), 300 µL of 100% ethanol was added to the lysate and then transferred to a DNA column (DNeasy Blood and Tissue Kit, Qiagen) and washed with 350 µL of buffer AW1 (Qiagen). Next, 80 µL of proteinase K solution was added to the DNA column and incubated at room temperature for 5 min. The DNA column was washed with buffer AW1 and buffer AW2. DNA was eluted using 2 × 30 µL elution buffer (buffer EB, Qiagen) into two separate fresh 1.5 mL centrifuge tubes for temporary storage at 4°C until processed for sequencing or in −80°C freezer for sample archiving.

#### *Sequence Processing*

We analyzed the sequence data using MOTHUR V.1.38.0 (seed = 777; Schloss et al., 2009) based on the MiSeq standard operating procedure accessed on 3 November 2015 and modified with time (see data accessibility and supplemental methods). Briefly, paired-end reads were merged into contigs based on the Phred quality score heuristic with *make.contigs* (Kozich et al., 2013). Contigs were filtered based on ambiguous bases, more than 8 homopolymers, a length outside of 240-275. Next, sequences were de-replicated and representative sequences were aligned with the Silva database and those not corresponding to the V4 region were removed. After the sequences were de-replicated and filtered, sequencing errors were removed using *pre.cluster* and chimeras were removed with UCHIME (Edgar et al., 2011). We then clustered representative sequences into OTUs at 97% similarity using the average neighbor algorithm (*cluster.split* command) and assigned the taxonomy of the OTUs using the Wang method implemented in the Ribosomal Database Project classifier (*classify.seqs* command).

#### *Sensitivity Analysis of Rare Taxa*

Due to the long tails within microbial rank-abundance curves (i.e. many rare taxa), we performed a sensitivity analysis to see the impact of rare taxa on the BEF relationships we observed (Figure S6). We removed OTUs that had a count of 1, 5, 10, 20, 30, 60, 90, 150, 225, and 300 sequences throughout the entire dataset and then checked the relationship with diversity versus community-wide and per-capita heterotrophic production (Figure S6). For example, removing 10-tons will be removing any OTUs that have a count of less than 10 sequences throughout the entire dataset. All code is for this supplementary analysis is available at: https://deneflab.github.io/Diversity_Productivity/analysis/OTU_Removal_Analysis.html

***Calculating Phylogenetic Diversity***

Instead of using Faith's phylogenetic diversity (Faith, 1992), we used the mean pairwise phylogenetic distance (or MPD), which measures the average phylogenetic distance between all combinations of two taxa pulled from the observed community. The MPD of the observed community was compared to the MPD of a null community with the same OTU richness and abundances randomized across OTUs pulled from all of the samples in the dataset. The difference between the observed MPD metric and randomized MPD metric were compared to each other while dividing by the standard deviation of the null community, known as the standardized effect size or SES (Gurevitch et al., 1993) of the MPD, or $SES_{MPD}$, (*ses.mpd* function in picante using null.model = "independentswap"; equation 1). We calculated both the abundance-unweighted and -weighted metrics of $SES_{MPD}$ (Webb et al., 2002). The $SES_{MPD}$ of each local community was calculated as follows:

$$SES_{MPD} = \frac{MPD_{Observed} - Mean(MPD_{Randomized})}{SD(MPD_{Randomized})} \qquad (1)$$ *(Kembel, 2009)*

Specifically, this model tests whether the mean $SES_{MPD}$ across samples differs from the null community (randomly generated from all samples to generate a randomized regional species pool) with an SES value of zero. Therefore, values higher than zero indicate phylogenetic evenness or overdispersion (higher phylogenetic diversity) while values less than zero indicate phylogenetic clustering (lower diversity) or that species are more closely related than expected according to the null community (Kembel, 2009).

#### *Standard Statistical Testing*

Data analysis was performed using R version 3.4.2 (R Core Team 2017), specifically with the phyloseq (McMurdie and Holmes, 2013), *stats* (R Core Team 2017), and *broom* (Robinson, 2017) R packages. All main figures were made using the ggplot2 R package (Wickham, 2009).

To assess a statistical difference in particle-associated and free-living cell abundances, community production rates, per-capita production rates, and biodiversity metrics, a Wilcoxon rank sum test (*wilcox.test* function) was performed. We evaluated whether diversity metrics or environmental variables predicted heterotrophic production rates using ordinary least squares linear regression (*lm* function) and accessed specific variables with *broom::glance()* (*i.e.* AIC, adjusted $R_2$). P-values were corrected using the false discovery rate method using the *p.adjust(method = "fdr")* function in the stats package.

### References


Edgar, R. C., B. J. Haas, J. C. Clemente, C. Quince, and R. Knight. 2011. UCHIME improves sensitivity and speed of chimera detection. Bioinformatics 27:2194–2200.

Faith, D. P. 1992. Conservation evaluation and phylogenetic diversity. Biological Conservation 61:1–10.

Fox, J., and S. Weisberg (2011). An {R} Companion to Applied Regression, Second Edition. Thousand Oaks CA: Sage. http://socserv.socsci.mcmaster.ca/jfox/Books/Companion

Gurevitch, J., L. L. Morrow, A. Wallace, and J. S. Walsh. 1992. A Meta-Analysis of Competition in Field Experiments. The American Naturalist 140:539–572.

Kembel, S. W. 2009. Disentangling niche and neutral influences on community assembly: Assessing the performance of community phylogenetic structure tests. Ecology Letters 12:949–960.

Kozich, J. J., S. L. Westcott, N. T. Baxter, S. K. Highlander, and P. D. Schloss. 2013. Development of a dual-index sequencing strategy and curation pipeline for analyzing amplicon sequence data on the MiSeq Illumina sequencing platform. Applied and Environmental Microbiology 79:5112–5120.

McMurdie, P. J., and S. Holmes. 2013. phyloseq: An R Package for Reproducible Interactive Analysis and Graphics of Microbiome Census Data. PLoS ONE 8:e61217.

R Core Team (2017) R: A Language and Environment for Statistical computing. Vienna, Austria: R Foundation for Statistical Computing. https:// www.R-project.org/.

Robinson, R. 2017. broom: Convert Statistical Analysis Objects into Tidy Data Frames. R package version 0.4.2. https://CRAN.R-project.org/package=broom

Schloss, P. D. et al. 2009. Introducing mothur: Open-source, platform-independent, community-supported software for describing and comparing microbial communities. Applied and Environmental Microbiology 75:7537–7541.

Webb, C. O., D. D. Ackerly, M. A. McPeek, and M. J. Donoghue. 2002. Phylogenies and Community Ecology. Annual Review of Ecology and Systematics 33:475–505.

Wickham, H. 2009. ggplot2: Elegant Graphics for Data Analysis. Springer-Verlag New York.

**Table S1**

Ordinary least squares linear regressions predicting **community-wide heterotrophic production** from all diversity and environmental variables with a FDR-corrected p-value of less than 0.05, sorted by AIC.

| fraction | independent_var | AIC | adj.r.squared | FDR.p.value |
|---|---|---|---|---|
| Particle | Inverse_Simpson | 74.34 | 0.69 | 0.0019 |
| Particle | Richness | 78.68 | 0.56 | 0.0063 |
| Particle | Shannon_Entropy | 79.61 | 0.52 | 0.0063 |
| Particle | Simpsons_Evenness | 80.85 | 0.47 | 0.0082 |
| All Samples | pH | 192.16 | 0.35 | 0.0329 |

**Table S2**

Ordinary least squares linear regressions predicting **per-capita heterotrophic production** from all diversity and environmental variables with a FDR-corrected p-value of less than 0.05, sorted by AIC.

| fraction | independent_var | AIC | adj.r.squared | FDR.p.value |
|---|---|---|---|---|
| Free | pH | -2.39 | 0.78 | 0.0025 |
| Particle | Inverse_Simpson | 8.29 | 0.69 | 0.0038 |
| Particle | Richness | 11.95 | 0.57 | 0.0075 |
| Particle | Shannon_Entropy | 12.43 | 0.55 | 0.0075 |
| Particle | Simpsons_Evenness | 13.65 | 0.49 | 0.0096 |
| All Samples | Richness | 24.72 | 0.63 | 0 |
| All Samples | Shannon_Entropy | 30.87 | 0.52 | 0.0001 |
| All Samples | Inverse_Simpson | 33.42 | 0.46 | 0.0003 |
| All Samples | unweighted_PD | 35.21 | 0.42 | 0.0124 |

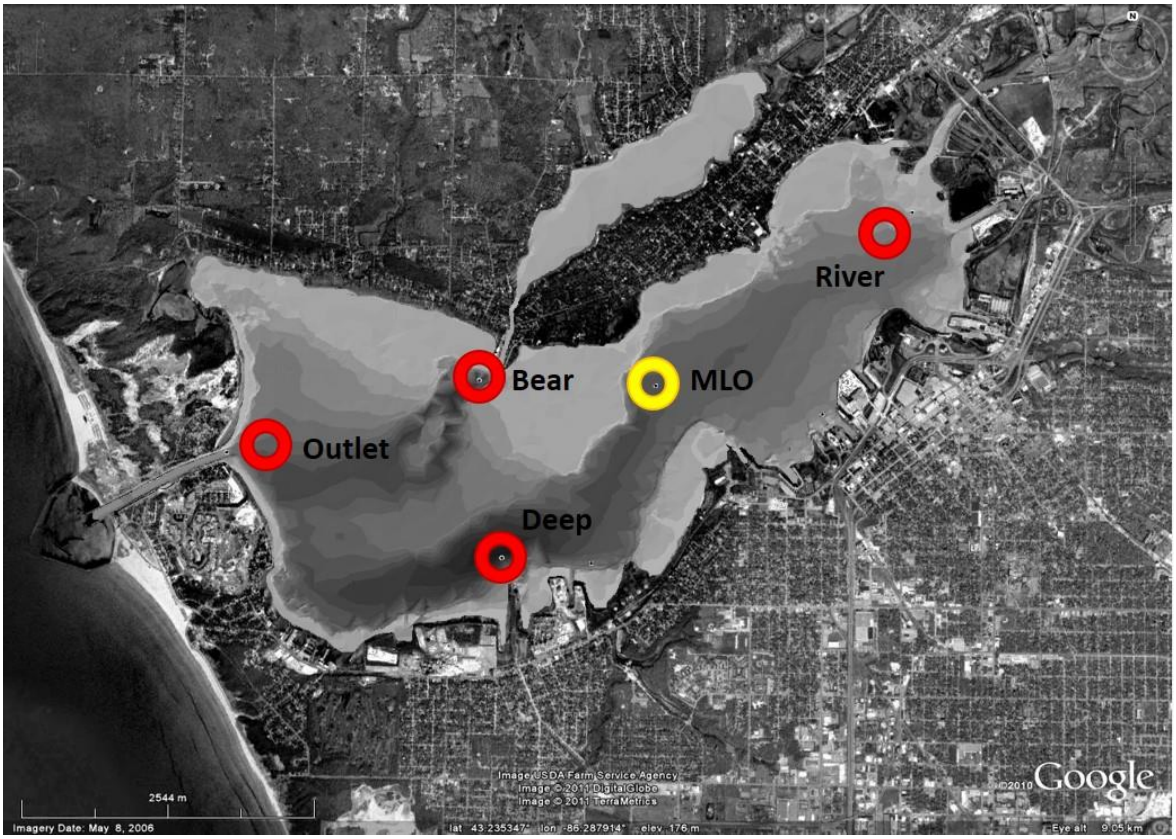


**Figure S1**

Bathymetric map of Muskegon Lake with locations of the Muskegon Lake Observatory Buoy (MLO) and the four sampling locations used in this study. Bathymetric iso-lines represent approximately 2m changes in water depth.

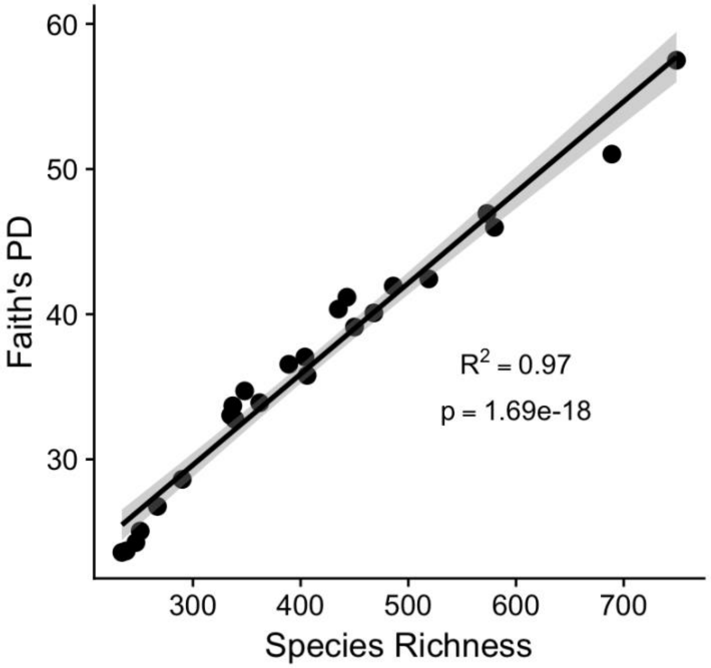


**Figure S2**

Faith's phylogenetic diversity is highly correlated with species richness and thus, it is important to compare standardized effect sizes that are measured when actual samples are compared to a randomized null model.

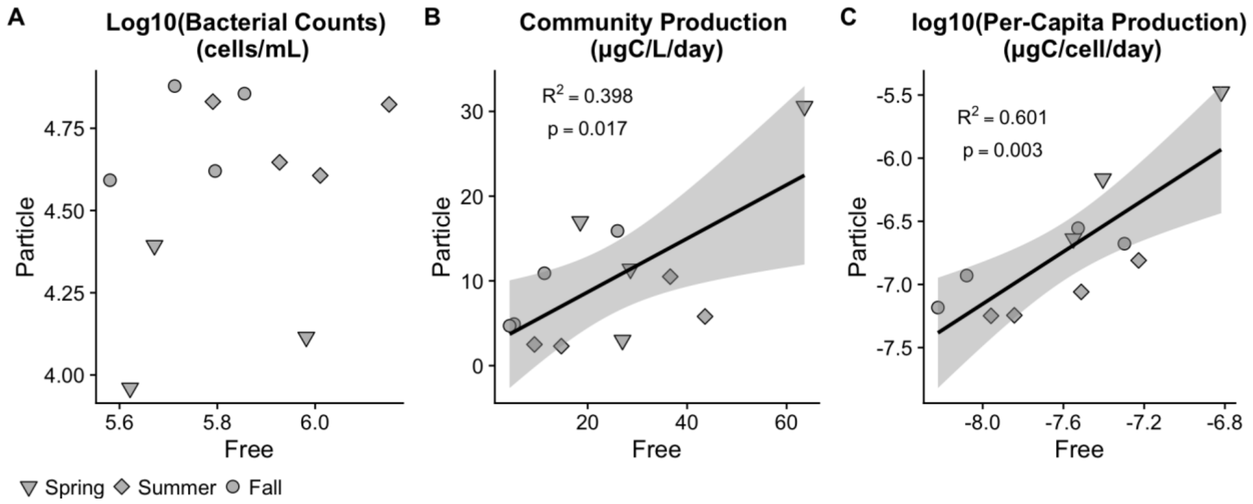


**Figure S3**

Correlation between corresponding particle-associated and free-living samples: **(A)** cell abundances in $log_{10}$(cells/mL), **(B)** community-wide heterotrophic production (ug C/L/day), **(C)** $log_{10}$**(**per-capita production) in ugC/cell/day.

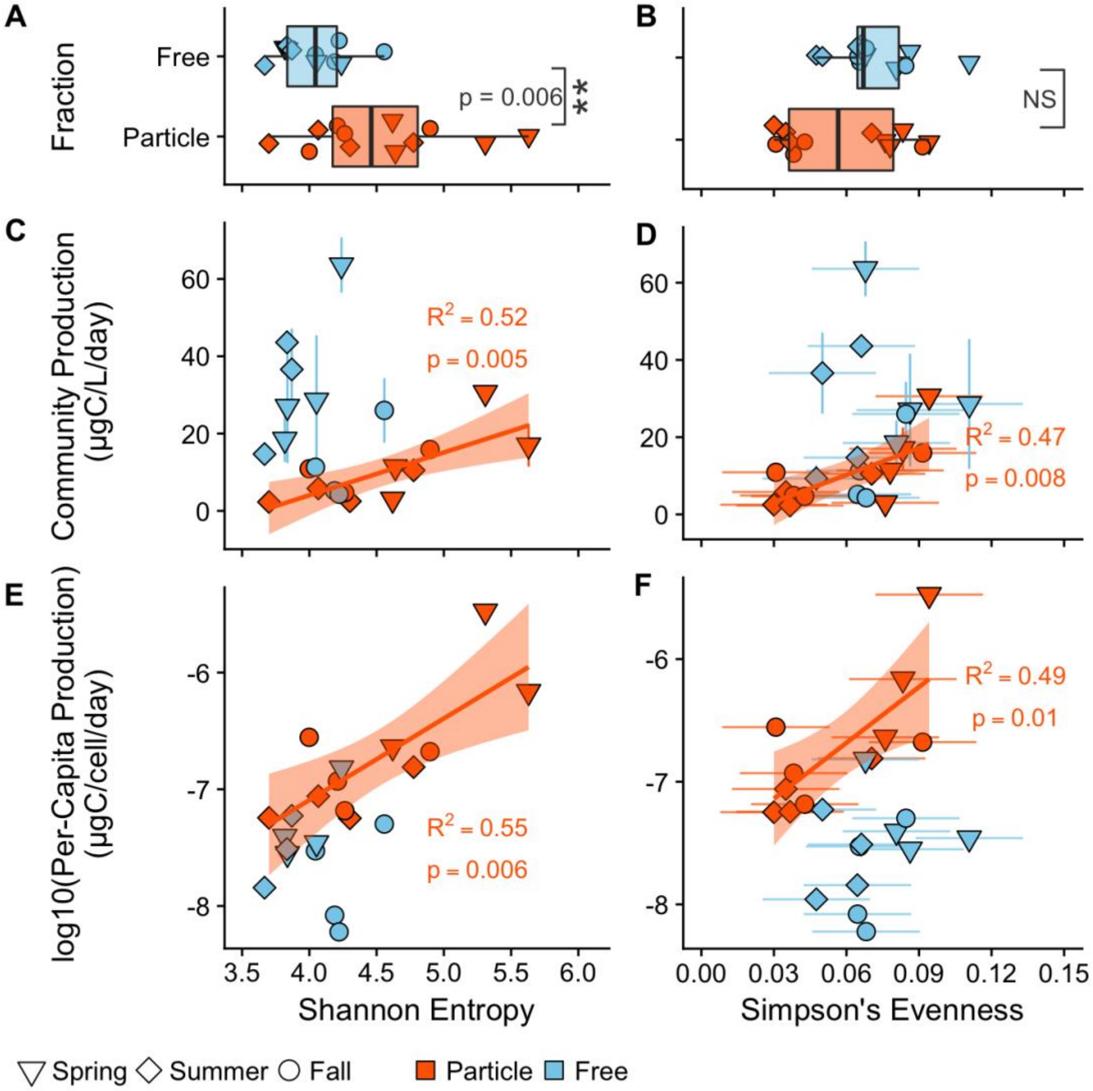


**Figure S4**

**Top Panel: (A)** Shannon entropy and **(B)** Simpson's evenness diversity metric between free-living (blue) and particle-associated (orange) habitats. **Middle panel:** The relationship between bacterial diversity and community-wide heterotrophic production (ugC/L/day). The y-axis is the same however, the x-axis represents **(C)** Shannon entropy and **(D)** Simpson's evenness. **Bottom panel:** The relationship between bacterial diversity and $\log_{10}$(heterotrophic production/cell) (ugC/cell/day). The y-axis is the same however, the x-axis represents **(E)** Shannon entropy and **(F)** Simpson's evenness. $R_2$ and p-values represented in the figures are outcomes of an ordinary least squares regression for the particle associated (orange) samples.

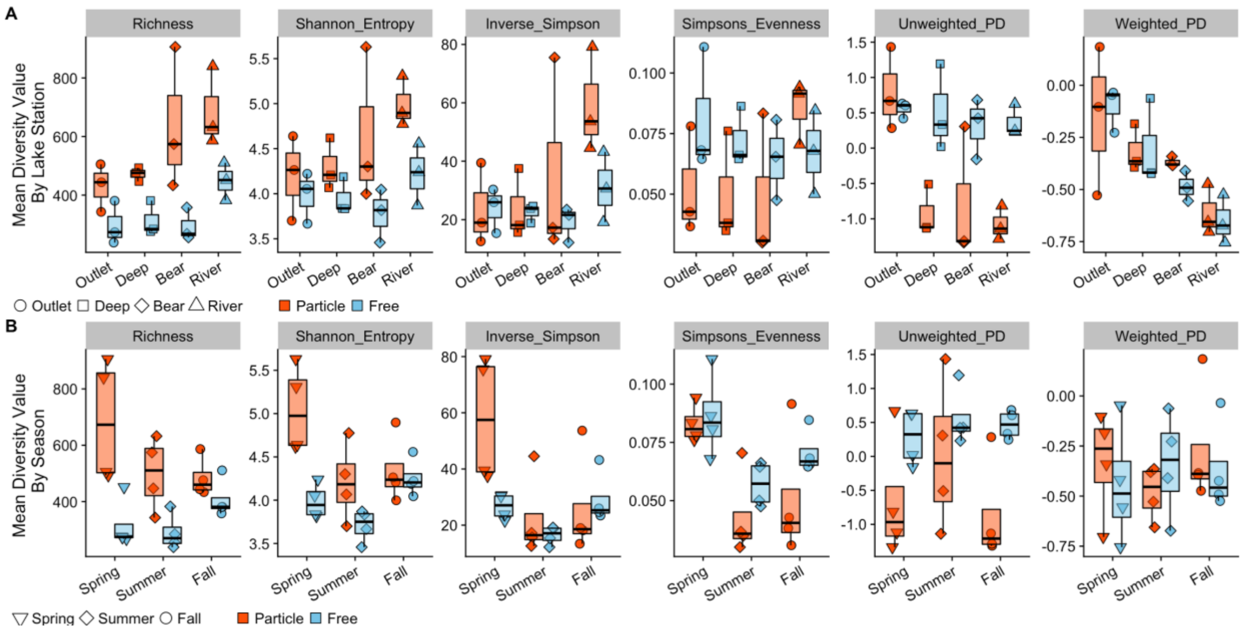


### Figure S5

**Top:** Particle-associated (orange) and free-living (blue) diversity values across the estuarine gradient in Muskegon Lake by station (from west to east). All diversity metrics calculated in this study are included. **Bottom:** Diversity values by season.

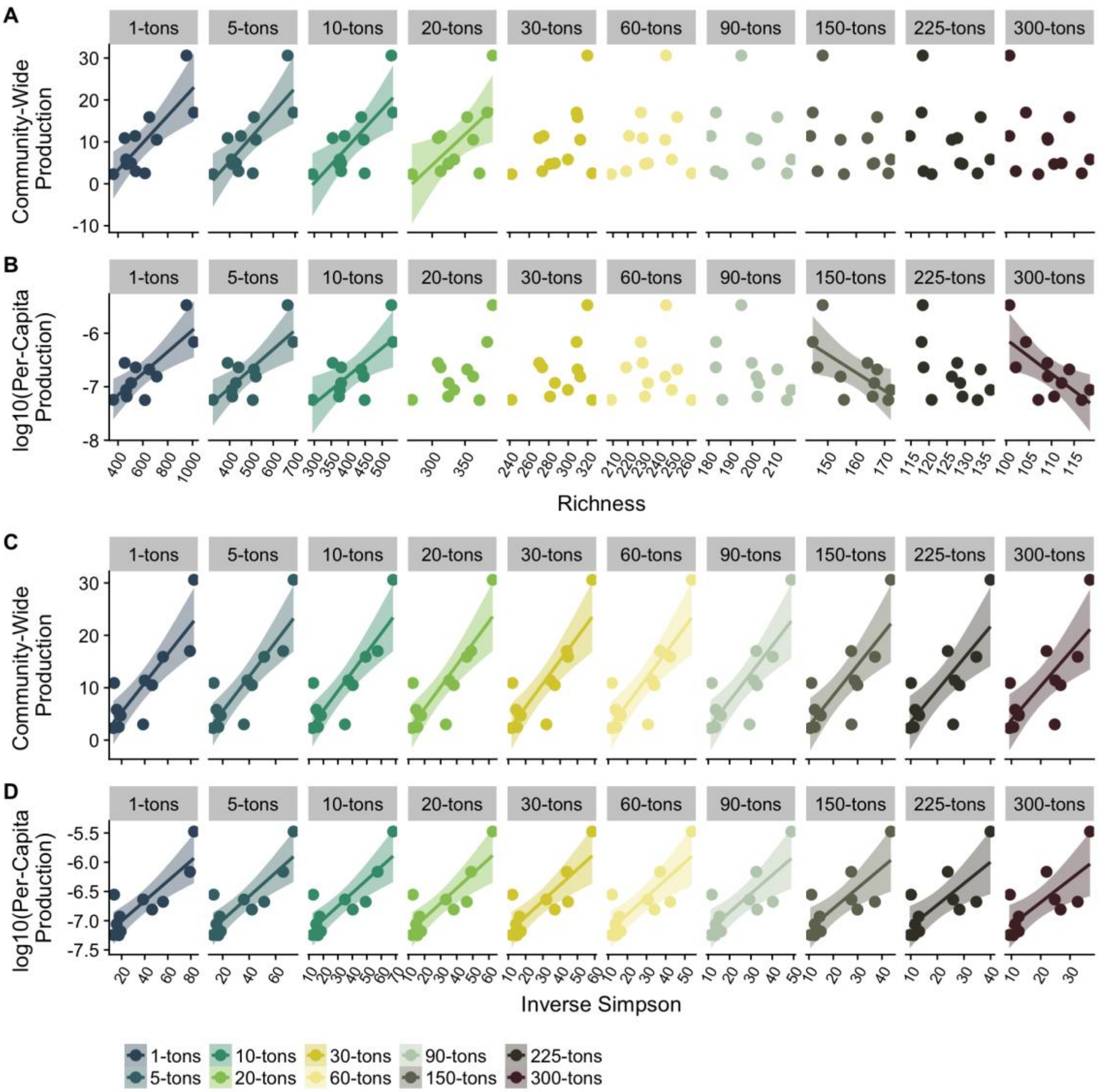


**Figure S6**

OTU removal analysis of particle-associated communities of singletons, doubletons, up to 300-tons. **(A)** and **(B)** represent the diversity-productivity patterns of observed richness while **(C)** and **(D)** represent the diversity-productivity relationship with the inverse Simpson's index. Plots **(A)** and **(C)** are community-wide heterotrophic production, while plots **(B)** and **(D)** are the $\log_{10}$(Per-capita heterotrophic production).

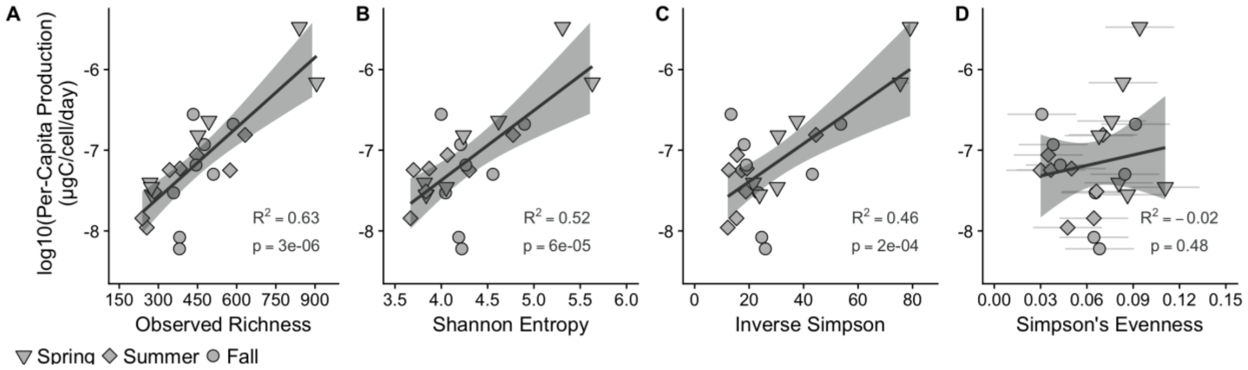


**Figure S7**

The relationship between bacterial diversity and $\log_{10}$(per-capita heterotrophic production) (ugC/cell/day) across all samples in the dataset. Diversity metrics on the x-axis are: **(A)** Observed richness, **(B)** Shannon entropy, **(C)** inverse Simpson's and **(D)** Simpson's evenness. Adjusted $R_2$ and p-values represent were calculated from an ordinary least squares linear regression model.

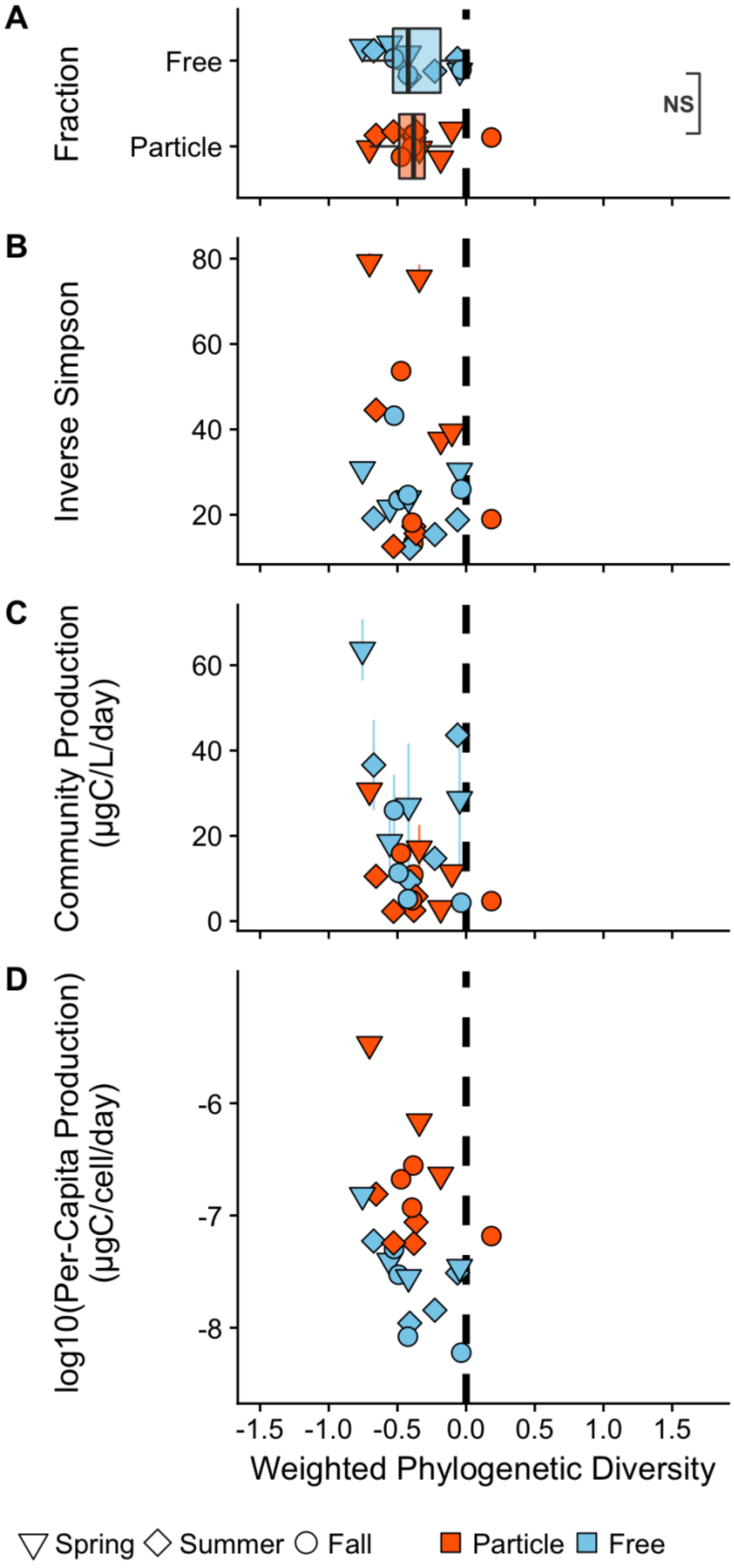


**Figure S8**

Abundance-weighted phylogenetic diversity **(A)** between particle-associated and free-living communities. **(B)** Absence of a correlation between abundance-weighted phylogenetic diversity and inverse Simpson's index, **(C)** community-wide heterotrophic production, and **(D)** per-capita production.

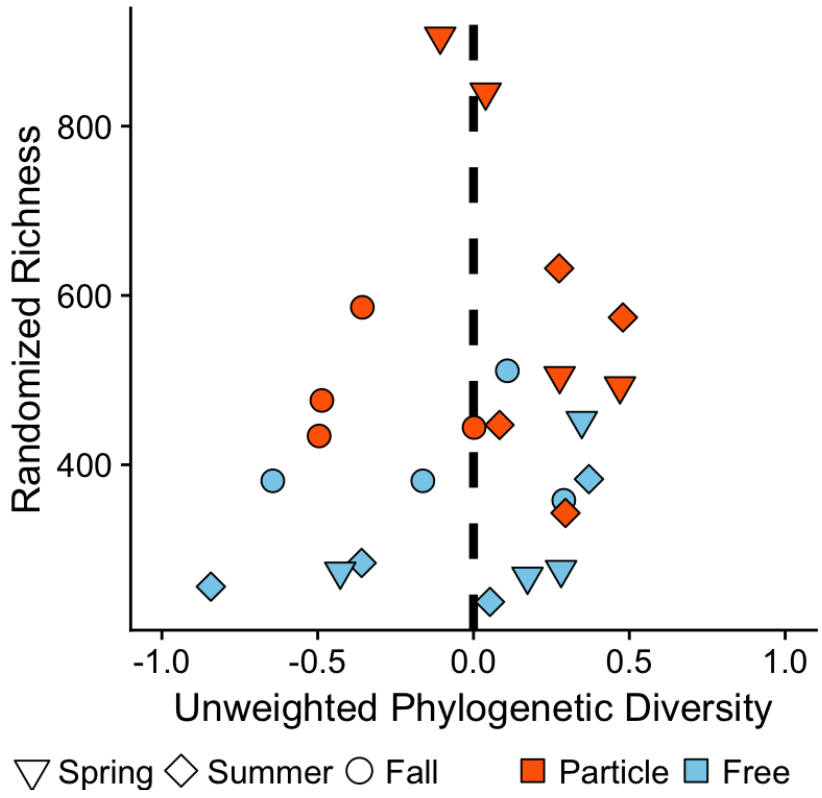


**Figure S9**

The relationship between randomized richness and standardized effect size. The richness values were the same value as actual samples, however, the OTUs across the samples were randomized across the gamma diversity of the entire dataset.

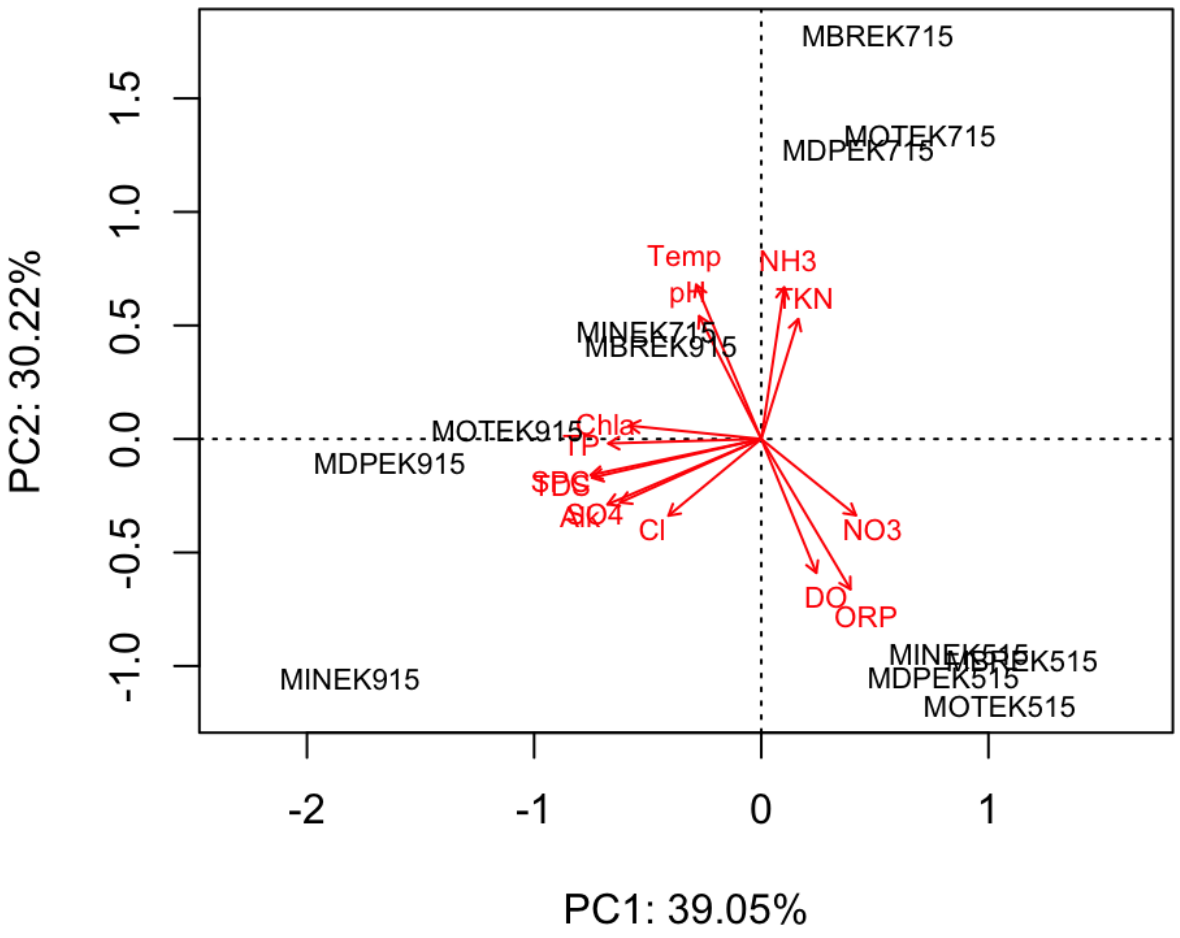


**Figure S10**

Principal components analysis of the euclidean distances of the environmental variables with a biplot (vectors) of the environmental drivers of stations in ordination space.
